# Neurofeedback training of auditory selective attention enhances speech-in-noise perception

**DOI:** 10.1101/2020.07.04.188045

**Authors:** Subong Kim, Caroline Emory, Inyong Choi

**Author notes:** Corresponding author at: Department of Communication Sciences and Disorders, University of Iowa, 250 Hawkins Dr., Iowa City, IA 52242, USA.

## Abstract

Selective attention enhances cortical responses to attended sensory inputs while suppressing others, which can be an effective strategy for speech-in-noise (SiN) understanding. Here, we introduce a training paradigm designed to reinforce attentional modulation of auditory evoked responses. Subjects attended one of two speech streams while our EEG-based attention decoder provided online feedback. After four weeks of this neurofeedback training, subjects exhibited enhanced cortical response to target speech and improved performance during a SiN task. Such training effects were not found in the Placebo group that underwent attention training without neurofeedback. These results suggest an effective rehabilitation for SiN deficits.

## Introduction

Albeit it is crucial for effective communication, the ability to understand speech in noise (SiN) differs dramatically even across normal-hearing individuals (Kumar *et al.*, 2007; Moore *et al.*, 2013). One reason for poor SiN understanding can be the deteriorated selective attention (Bressler *et al.*, 2017), as past studies showed a correlation between selective attention and SiN performance (Strait and Kraus, 2011). Compared to poor performers, listeners with good SiN performance exhibit greater amplitude ratio of cortical responses to target speech compared to responses to noise (Kim *et al.*, 2019), which may indicate that attentional modulation on neural encoding of acoustic inputs in the auditory cortex (AC) (Hillyard *et al.*, 1973; Mesgarani and Chang, 2012; Carcea *et al.*, 2017) is a key neural mechanism for successful SiN understanding.

Conventional hearing remediations through amplification cannot improve SiN understanding ability (Bentler *et al.*, 2008). Instead, perceptual training is often considered as a solution for SiN difficulties (Whitton *et al.*, 2014; Whitton *et al.*, 2017). However, a frequently reported problem of perceptual training is that the training effect does not generalize to other auditory stimuli not used for the training (Fiorentini and Berardi, 1981; Wright *et al.*, 1997). This generalization problem leads us to consider a training that directly improves a key strategy for the SiN understanding: a training that reinforces attentional modulation of auditory cortical responses.

How, then, can we design a perceptual training paradigm that aims to reinforce attentional modulation of cortical activity? Theories of learning claim that the target of training is manipulated by rewarding; the determination of feedback (i.e., reward or punishment) must be based on the target training component (Goodman and Wood, 2004). Thus, to enhance attentional modulation of cortical responses, a training paradigm should provide feedback based on the strength of attentional modulation. Auditory selective attention can be reliably decoded from single-trial electroencephalographic (EEG) signals (Kerlin *et al.*, 2010; Choi *et al.*, 2013; O’Sullivan *et al.*, 2015). Since the accuracy of attention decoding from single-trial EEG signals reflects the strength of attentional modulation on cortical auditory evoked responses (Choi *et al.*, 2013), providing the result of EEG-based attention decoding as neurofeedback (Sherlin *et al.*, 2011) may reinforce the users’ attentional modulation of cortical responses. The goal of the present study is to provide evidence to support the concept of auditory selective attention training through such an EEG-based neurofeedback paradigm and explore its efficacy for SiN understanding ability.

## Methods

### Participants

Twenty young adult subjects with normal hearing, who were native speakers of American English, were recruited for this study (Mean age = 23.2 years; SD = 1.33 years; 6 (30%) male). Upon agreeing to the study, subjects were randomly assigned to either the Experimental or the Placebo (control) group (i.e., single-blinded design). All subjects completed four consecutive weeks of one-hour-per-week training and pre- and post-training SiN tests at their first and last visits. All study procedures were reviewed and approved by the University of Iowa Institutional Review Board.

### Experimental design and procedures

#### Attention training procedure: Experimental group

During each training session, three overlapping auditory streams were presented; 1) a male voice saying the word “down” repeated four times from the right (+30° azimuth) loudspeaker, 2) a female voice saying the word “up” repeated five times from the left (−30°) loudspeaker, 3) and a distractor non-speech noise that sounds like water bubble played intermittently from the loudspeaker directly in front of the subject. For each of the 120 trials in each visit, a visual cue (“Target: Up” or “Target: Down”) was given to direct participants’ attention to either “up” or “down” stream (60 trials each). After sounds, listeners’ attention was decoded from EEG. A visual feedback (“+” sign on the screen moving up or down) was given at the end of a trial to indicate the decoded direction of attention (i.e., attended “up” or “down” stream, respectively). **Figure 1** illustrates an example of a trial attending the “down” stream.

**Figure 1.**
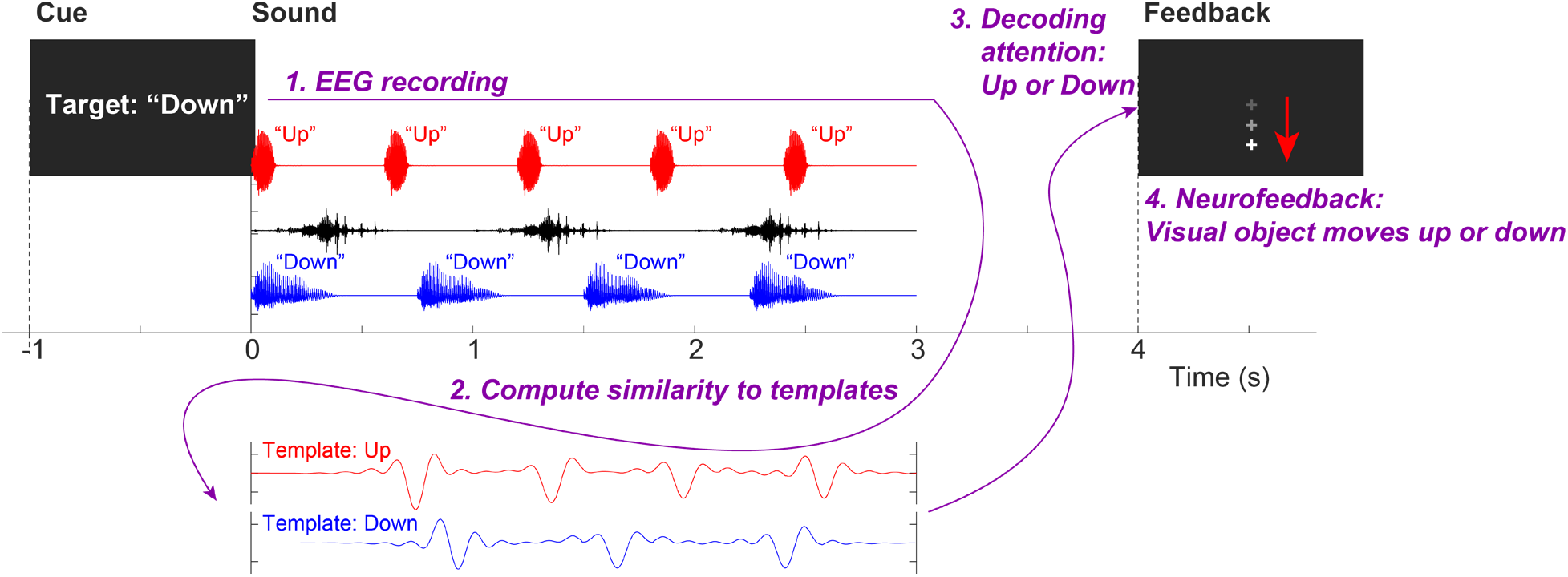
Trial structure of the neurofeedback training assigned to the Experimental group. This example shows an attend-*down* trial.

#### Attention training procedure: Placebo group

The Placebo group listened to the similar three overlapping auditory streams (i.e., isochronous repetitions of “up” and “down” spoken by the female and male speakers) where one of the last 3 or 2 utterances in each stream had 3-semi-tone higher pitch. As the visual cue directed their attention to either “up” or “down” stream in each trial, they picked an utterance with a higher pitch in the attended stream by pressing number key (i.e., an oddball detection task). After the button press, visual feedback (“Correct” or “Incorrect”) was given based on the accuracy of their button response.

#### Pre- and post-training speech-in-noise tests

All subjects, regardless of group designation, completed the same pre- and post-training SiN test. The test used 100 monosyllabic consonant-vowel-consonant English words from a pre-recorded California Consonant Test with added multi-talker babble noise. Stimuli were presented at ±3 dB SNR (50 words each) by changing the level of the noise in random order. At each trial, a target word started 1 second after the noise onset. At the end of a trial, subjects picked a word they heard from four choices given on the screen. For further analysis, behavioral and neural data from −3 dB SNR condition showing larger individual differences and no ceiling effect were used.

### ERP analysis

Sixty-four channel scalp EEG data were recorded during the training and SiN tasks using the BioSemi ActiveTwo system at a 2048 Hz sampling rate with the international 10-20 configuration. In order to provide neurofeedback to the Experimental group, a template-matching method was used to decode attention from single-trial EEG signals (Choi *et al.*, 2013). EEG recordings from front-central channels (Fz, FCz, FC1, FC2, Cz) were averaged and re-referenced to linked mastoids. EEG signals were bandpass-filtered between 1 and 9 Hz, baseline corrected, and then compared to two pre-generated template EEG waveforms obtained from cortical evoked responses to “up” and “down” streams. The attention was decoded by finding the template that has a larger correlation coefficient with the EEG signal. For analyzing EEG data obtained during SiN tasks, after applying a bandpass filter between 1 and 30 Hz using a 2048-point FIR filter, epochs were extracted and baseline-corrected. After correcting ocular artifacts using independent component analysis (Jung *et al.*, 2000), the epochs were averaged at each electrode. Since we use non-repeating naturally spoken words as stimuli, the latency of event-related potentials (e.g., N1) varied across words. To obtain clean N1 from averaged evoked response, every epoch was rearranged according to the median N1 latency of its corresponding word obtained from the grand mean of 50 normal hearing subjects who completed the same SiN task in our laboratory previously.

In order to project the sensor-space data into source-space, the inverse operator was estimated using minimum norm estimation (MNE) (Hämäläinen, 1989; Gramfort *et al.*, 2013; Gramfort *et al.*, 2014) based on assumptions of multiple sparse priors (Friston *et al.*, 2008) on an average template brain. To focus on non-speech-specific attention effect on AC while avoiding confounding factors related to language processing, we chose right – instead of left – Heschl’s gyrus (HG) as the region of interest.

### Statistical analysis

Two-way mixed ANOVAs were conducted on both behavioral performance and neural data. To get clear ERPs and perform statistical analysis on neural data, we computed leave-one-out grand averages (i.e., jackknife approach). In addition, ERP envelopes were extracted by applying a 4 – 8 Hz bandpass filter and Hilbert transform for ERP magnitude comparisons. The peak magnitudes of ERP envelopes obtained at ~230 ms after the word onset from the right HG were compared between conditions. As post hoc analysis, one-tailed *t*-tests were conducted with Bonferroni correction as we were only interested in the improvement (i.e., one direction) in the behavioral performance (accuracy) and acoustic encoding at AC after training. Inflated *F*-ratios and *t*-values due to the jackknife approach were adjusted (Luck, 2014).

## Results

### Behavioral performance

We found that the behavioral performance (accuracy) did not improve after training [*F*_1,18_ = 3.37, *p* = 0.083] on average; however, accuracy did increase significantly in the Experimental group [*t*_9_ = −2.37, *p* = 0.042], but not in the Placebo group [*t*_9_ = −0.71, *p* = 0.49] (**Figure 2A**). We found no group effect [*F*_1,18_ = 2.49, *p* = 0.13] on average; however, the post hoc analysis showed a significant difference in performance between two groups after training [*t*_18_ = 2.15, *p* = 0.045], but not before training [*t*_18_ = 0.60, *p* = 0.56] (**Figure 2A**). No interaction between training and group [*F*_1,18_ = 0.38, *p* = 0.55] was revealed.

**Figure 2.**
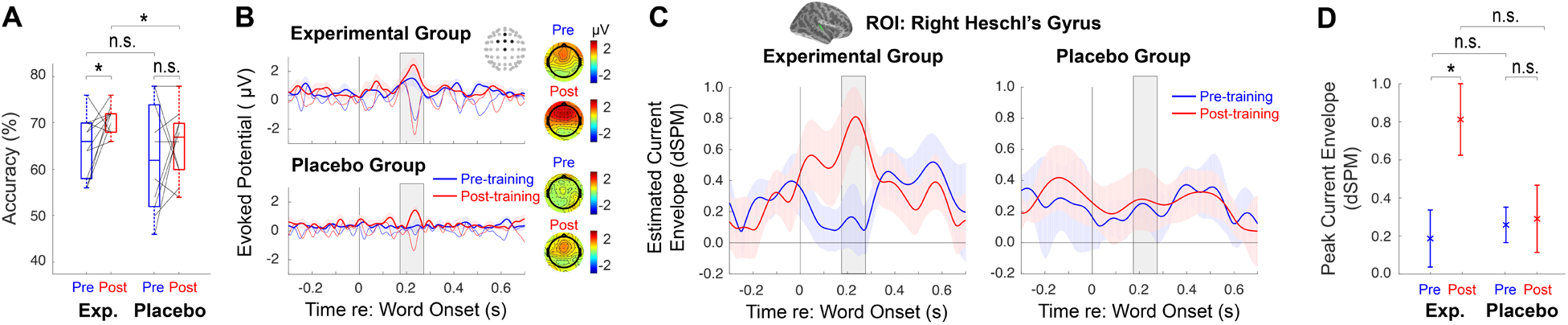
Training effects on **A**. accuracy, **B**. sensor-space evoked responses (thick lines: envelopes) and topographies, **C**. source time courses (current envelopes with ±1 standard error) in the right HG, and **D**. the peak evoked magnitude in the right HG. A significant training effect on the peak evoked magnitude is found in the Experimental group.

### Attention during training

During the attention training, both Experimental and Placebo groups showed a significant effect of attention on evoked responses as reflected in the single-trial attention decoding accuracy being higher than chance level (one-sample *t*-test, Experimental group: *t*_*9*_ = 8.34, *p* < 0.001, Placebo group: *t*_*9*_ = 5.12, *p* < 0.001). The mean accuracy of single-trial attention decoding was 59.1% and 58.1%, with a standard deviation of 3.5% and 5.0% for the Experimental and Placebo groups, respectively. There was no significant difference in the single-trial classification accuracies between the groups (two-sample t-test, *p* = 0.59). The mean accuracy of the oddball detection in the Placebo group was 98.8%, with a 1.4% standard deviation.

### Source-space ERPs to SiN

Source-space data obtained from the right hemisphere HG showed that the AC response to SiN increased significantly after training [*F*_1,18_ = 4.78, *p* = 0.042]; notably, the training effect only appeared in the Experimental group [*t*_9_ = −3.16, *p* = 0.023], but not in the Placebo group [*t*_9_ = −0.14, *p* = 1] (**Figure 2D**). The *F*-test results revealed no group effect [*F*_1,18_ = 1.94, *p* = 0.18] on average; the post hoc analysis showed no significant difference in AC response to SiN between two groups before [*t*_18_ = −0.41, *p* = 1] and after training [*t*_18_ = 2.024, *p* = 0.12] (**Figure 2D**). There existed no significant interaction between training and group [*F*_1,18_ = 3.89, *p* = 0.064].

## Discussion

This study demonstrated that the neurofeedback training of auditory selective attention is effective for improving neural encoding and accurate speech recognition in background noise. As it is evident that attentional gain control process (Hillyard *et al.*, 1998) is involved in a successful speech-in-noise perception (Mesgarani and Chang, 2012; Kim *et al.*, 2019), our training paradigm was designed to reinforce attentional modulation of auditory cortical evoked potentials by providing visual feedback determined by an EEG-based attention decoder. After four weeks of training, we found consistent improvement in listeners’ SiN performance and acoustic encoding at AC only in the Experimental group.

Better representation of target speech at AC may reflect an active sensory gain control for the Experimental group after training (Shinn-Cunningham and Best, 2008). This is consistent with the previous finding that showed attention could modulate the sound representation in AC and improve behavioral performance (Mesgarani and Chang, 2012; Carcea *et al.*, 2017). Attention may increase the gain at the neural population level by increasing the response of neurons to the target word. Learning or training can improve selective enhancement of neural response, that may develop over time and last longer, and improve speech perception (Froemke *et al.*, 2013). The training effect found in the present study may develop and last over weeks and result in an improvement in speech perception.

Our Experimental group training was differentiated from the Placebo training in that the feedback reward was determined by participants’ neural activity, not by behavioral performance. Similarly to the reports by Whitton *et al.* (2014) and Whitton *et al.* (2017), the present study showed that the effect of neurofeedback training was transferred to SiN performance, while the Placebo group did not show such generalizability. The generalizability of the training effect observed in the present study may indicate that the reinforcement of attentional modulation on cortical responses may improve a key neural strategy for the SiN understanding. In contrast, our Placebo training provided a primary task of detecting pitch oddballs from the attended stream. Feedbacks provided to the Placebo group informed how their oddball detection was accurate, not how strong their attention was. Our results may indicate the “indirect” primary task that does not provide a reward shaped by the strength of attention would not exhibit generalizability of its training effect to the speech in noise task.

## Acknowledgments

This work was supported by American Otological Society Research Grant and Department of Defense Hearing Restoration Research Program Grant (W81XWH-19-1-0637) awarded to Inyong Choi, as well as NIDCD P50 (DC000242 31). The authors declare no conflicting financial interests.

## References

Bentler, R., Wu, Y. H., Kettel, J., and Hurtig, R. (2008). “Digital noise reduction: outcomes from laboratory and field studies,” International journal of audiology 47, 447–460.

Bressler, S., Goldberg, H., and Shinn-Cunningham, B. (2017). “Sensory coding and cognitive processing of sound in Veterans with blast exposure,” Hear Res 349, 98–110.

Carcea, I., Insanally, M. N., Froemke, R. C., Carcea, I., Insanally, M. N., and Froemke, R. C. (2017). “Dynamics of auditory cortical activity during behavioural engagement and auditory perception,” Nat. Commun. Nature Communications 8.

Choi, I., Rajaram, S., Varghese, L. A., and Shinn-Cunningham, B. G. (2013). “Quantifying attentional modulation of auditory-evoked cortical responses from single-trial electroencephalography,” Front Hum Neurosci 7, 115.

Fiorentini, A., and Berardi, N. (1981). “Learning in grating waveform discrimination: Specificity for orientation and spatial frequency,” VR Vision Research 21, 1149,1153–1151,1158.

Friston, K., Harrison, L., Daunizeau, J., Kiebel, S., Phillips, C., Trujillo-Barreto, N., Henson, R., Flandin, G., and Mattout, J. (2008). “Multiple sparse priors for the M/EEG inverse problem,” Neuroimage 39, 1104–1120.

Froemke, R. C., Carcea, I., Barker, A. J., Yuan, K., Seybold, B. A., Martins, A. R., Zaika, N., Bernstein, H., Wachs, M., Levis, P. A., Polley, D. B., Merzenich, M. M., and Schreiner, C. E. (2013). “Long-term modification of cortical synapses improves sensory perception,” Nat Neurosci 16, 79–88.

Goodman, J. S., and Wood, R. E. (2004). “Feedback specificity, learning opportunities, and learning,” The Journal of applied psychology 89, 809–821.

Gramfort, A., Luessi, M., Larson, E., Engemann, D. A., Strohmeier, D., Brodbeck, C., Goj, R., Jas, M., Brooks, T., Parkkonen, L., and Hamalainen, M. (2013). “MEG and EEG data analysis with MNE-Python,” Front Neurosci 7, 267.

Gramfort, A., Luessi, M., Larson, E., Engemann, D. A., Strohmeier, D., Brodbeck, C., Parkkonen, L., and Hamalainen, M. S. (2014). “MNE software for processing MEG and EEG data,” Neuroimage 86, 446–460.

Hämäläinen, M. S., Sarvas, J. (1989). “Realistic conductivity geometry model of the human head for interpretation of neuromagnetic data,” IEEE Trans Biomed Eng 36, 165–171.

Hillyard, S. A., Hink, R. F., Schwent, V. L., and Picton, T. W. (1973). “Electrical signs of selective attention in the human brain,” Science 182, 177–180.

Hillyard, S. A., Vogel, E. K., and Luck, S. J. (1998). “Sensory gain control (amplification) as a mechanism of selective attention: electrophysiological and neuroimaging evidence,” Philosophical transactions of the Royal Society of London. Series B, Biological sciences 353, 1257–1270.

Jung, T. P., Makeig, S., Humphries, C., Lee, T. W., McKeown, M. J., Iragui, V., and Sejnowski, T. J. (2000). “Removing electroencephalographic artifacts by blind source separation,” Psychophysiology 37, 163–178.

Kerlin, J. R., Shahin, A. J., and Miller, L. M. (2010). “Attentional gain control of ongoing cortical speech representations in a “cocktail party”,” J Neurosci 30, 620–628.

Kim, S., Schwalje, A., Liu, A., Gander, P., McMurray, B., Griffiths, T., and Choi, I. (2019). “External and Internal Signal-to-noise Ratios Alter Timing and Location of Cortical Activities During Speech-in-noise Perception,” bioRxiv 817460.

Kumar, G., Amen, F., and Roy, D. (2007). “Normal hearing tests: is a further appointment really necessary?,” Journal of the Royal Society of Medicine Journal of the Royal Society of Medicine 100, 66.

Luck, S. J. (2014). An introduction to the event-related potential technique (The MIT Press, Cambridge, Massachusetts).

Mesgarani, N., and Chang, E. F. (2012). “Selective cortical representation of attended speaker in multi-talker speech perception,” Nature 485, 233–236.

Moore, D. R., Rosen, S., Bamiou, D.-E., Campbell, N. G., and Sirimanna, T. (2013). “Evolving concepts of developmental auditory processing disorder (APD): A British Society of Audiology APD Special Interest Group white paper,” International journal of audiology 52, 3–13.

O’Sullivan, J. A., Power, A. J., Mesgarani, N., Rajaram, S., Foxe, J. J., Shinn-Cunningham, B. G., Slaney, M., Shamma, S. A., and Lalor, E. C. (2015). “Attentional Selection in a Cocktail Party Environment Can Be Decoded from Single-Trial EEG,” Cerebral Cortex 25, 1697–1706.

Sherlin, L. H., Arns, M., Lubar, J., Heinrich, H., Kerson, C., Strehl, U., and Sterman, M. B. (2011). “Neurofeedback and Basic Learning Theory: Implications for Research and Practice,” Journal of Neurotherapy 15, 292–304.

Shinn-Cunningham, B. G., and Best, V. (2008). “Selective attention in normal and impaired hearing,” Trends Amplif 12, 283–299.

Strait, D. L., and Kraus, N. (2011). “Can you hear me now? Musical training shapes functional brain networks for selective auditory attention and hearing speech in noise,” Front Psychol 2, 113–113.

Whitton, J. P., Hancock, K. E., and Polley, D. B. (2014). “Immersive audiomotor game play enhances neural and perceptual salience of weak signals in noise,” procnatiacadscie Proceedings of the National Academy of Sciences of the United States of America 111, 9030.

Whitton, J. P., Hancock, K. E., Shannon, J. M., and Polley, D. B. (2017). “Audiomotor Perceptual Training Enhances Speech Intelligibility in Background Noise,” Current biology : CB. 27, 3237–3247.e3236.

Wright, B. A., Buonomano, D. V., Mahncke, H. W., and Merzenich, M. M. (1997). “Learning and generalization of auditory temporal-interval discrimination in humans,” The Journal of neuroscience : the official journal of the Society for Neuroscience 17, 3956–3963.

